# Beyond screens: a gaze-driven auditory human-computer interface

**DOI:** 10.64898/2026.11.11.765668

**Authors:** Prachi Patel, Vinay Raghavan, Xiaomin He, Nima Mesgarani

## Abstract

Brain-computer interfaces enable humans to communicate and manipulate external devices by converting physiological signals into actions. These signals often have a signal-to-noise ratio (SNR) too low to enable their use in real-world Human-Computer Interaction (HCI). Notably, electrooculography (EOG) measurements of gaze direction exhibit a higher SNR, making them widely employed in HCI applications with displays. Nevertheless, display-free paradigms can also leverage the high SNR of EOG measurements of eye gaze. Current auditory paradigms present options sequentially, extending stimulus presentation time and reducing the information transfer rates (ITRs) compared to visual displays. This paper introduces an HCI approach that combines the SNR of eye gaze with a rapid auditory display paradigm. This paradigm leverages the human capacity to focus on a single auditory stream presented among others. By integrating eye gaze with a rapid auditory display, our method attains average accuracies of 82.7% for a five-option menu and average peak ITRs of 41 bits/min. This increase in performance represents a stride towards practical HCIs in real-world scenarios and display-free smart devices.

## I. Introduction

Brain-Computer Interfaces (BCI) are an evolving technology enabling human communication and control over external devices through the translation of physiological signals into actionable commands [1]. As the prevalence of smart devices rises, there is a growing interest and expanded research focus on BCIs. The exploration of alternative methods for human-computer interaction (HCI) remains a key objective in current research, with a specific emphasis on identifying physiological signals that offer increased intuitiveness in execution, enhanced robustness in decoding, and reduced intrusiveness in measurement for BCI applications.

Various physiological signals have found application in BCIs to facilitate diverse methods of HCI. For instance, non-invasive electroencephalography (EEG) has been employed to decode a user’s intent through event-related potentials (ERPs) in a visual speller paradigm [2]. The exploration of auditory displays as a screen-free, portable HCI option has been pursued in situations where visual displays are impractical [3]. Additionally, the utilization of electrical signals from muscles (electromyography, EMG) has extended to controlling robotic arms or interfacing with personal computers [4]. The amalgamation of multiple physiological signals has been leveraged to enhance HCI performance [5], [6]. Originally developed to assist individuals with disabilities, these technologies have transcended their initial scope and found applications in gaming, education, and health, offering a user-friendly, intuitive, private, and robust alternative to traditional input devices like joysticks, mice, keyboards, or speech [4], [7], [8]. However, challenges such as the low signal-to-noise ratio (SNR) of physiological signals like EEG have hindered their widespread use in HCI.

Notably, eye gaze emerges as a physiological signal measurable through eye-tracking cameras or electrooculography (EOG) with a significantly higher SNR compared to signals like EEG. Eye gaze finds popularity in HCI applications involving displays, such as virtual and augmented reality, and has been employed for tasks like cursor control [9], option selection [10] and speller tasks [11]. Nevertheless, scenarios arise where the use of screens is not feasible. For instance, individuals wearing smart headphones or glasses may need to skip a song, decline a call during a meeting, send a message while driving, or interact with a smart device with privacy concerns. In such contexts, the integration of eye gaze’s high SNR measurement with a display-free interface emerges as a highly desirable solution.

Recent years have witnessed the emergence of HCI methods extending beyond visual displays. An illustrative instance is the exploration of auditory displays for screen-free HCI [12]. Within this domain, the Auditory MUlti-class Spatial ERP (AMUSE) paradigm has been introduced [13], involving the presentation of multiple auditory options to a listener who then focuses on the desired option. The selected option is subsequently decoded from the Event-Related Potentials (ERPs) in the electroencephalography (EEG) signals. These paradigms hold practicality, as they can be seamlessly integrated into everyday form factors like glasses or headphones. However, auditory HCI methods utilizing EEG exhibit lower performance, measuring around 10 bits/min [14], in contrast to visual HCI methods which achieve approximately 20 bits/min [15], assessed by the standard metric of Information Transfer Rate (ITR). This diminished performance is attributed to the low Signal-to-Noise Ratio (SNR) of EEG signals in auditory HCIs and the sequential nature of auditory displays. The latter extends the presentation time of options compared to visual displays that present all options simultaneously on a screen. To enhance ITR, our proposed approach leverages the human capacity to selectively attend to and isolate a single auditory stream when presented with multiple simultaneous streams. This is achieved by considering the spatial and spectrotemporal features of the target audio stream [16], [17], [18], [19]. By capitalizing on this inherent ability, it becomes feasible to modify the standard AMUSE paradigm, overlapping auditory options in time, and thereby reducing option presentation time This paper introduces an innovative approach to human-computer interaction by merging the elevated SNR of eye gaze, measured through EOG, with a modified AMUSE paradigm featuring a rapid, overlapped auditory display. The synergy of these elements results in a significantly heightened information transfer rate, paving the way for more efficient communication with smart devices.

## II. Methods

### A. Participants

Ten adults (6 females and 4 males) with an age range of 22-39 years and self-reported normal hearing participated in this study. Two additional participants took part in this experiment but were not included in the analysis due to poor data quality. The experiment was undertaken in accordance with the Declaration of Helsinki, and the Institutional Review Board of Columbia University approved this study. All participants were given a briefing on the study and provided written consent. They were also compensated for their time.

### B. Experiment

During the study, the participants were asked to sit in front of a computer monitor in a private room. The sounds were presented using standard Panasonic stereo in-ear headphones (Panasonic RP-HJE120) at a sampling rate of 16 kHz. The stimuli consisted of five target words– “one,” “two,” “three,” “four,” and “five”–presented from -60, -30, 0, 30 and 60 degrees in the azimuthal plane, respectively (Fig. 1A). The stimuli were exactly 550 milliseconds long and were spatialized using binaural room impulse response filters from github.com/IoSR-Surrey/RealRoomBRIRs. The sounds were synthesized using the Google Cloud Text-to-Speech API with three unique female voices for “one,” “three,” and “five,” and two unique male voices for “two” and “four.” The loudness of the stimulus was adjusted to a comfortable level for each participant.

**Fig 1.**
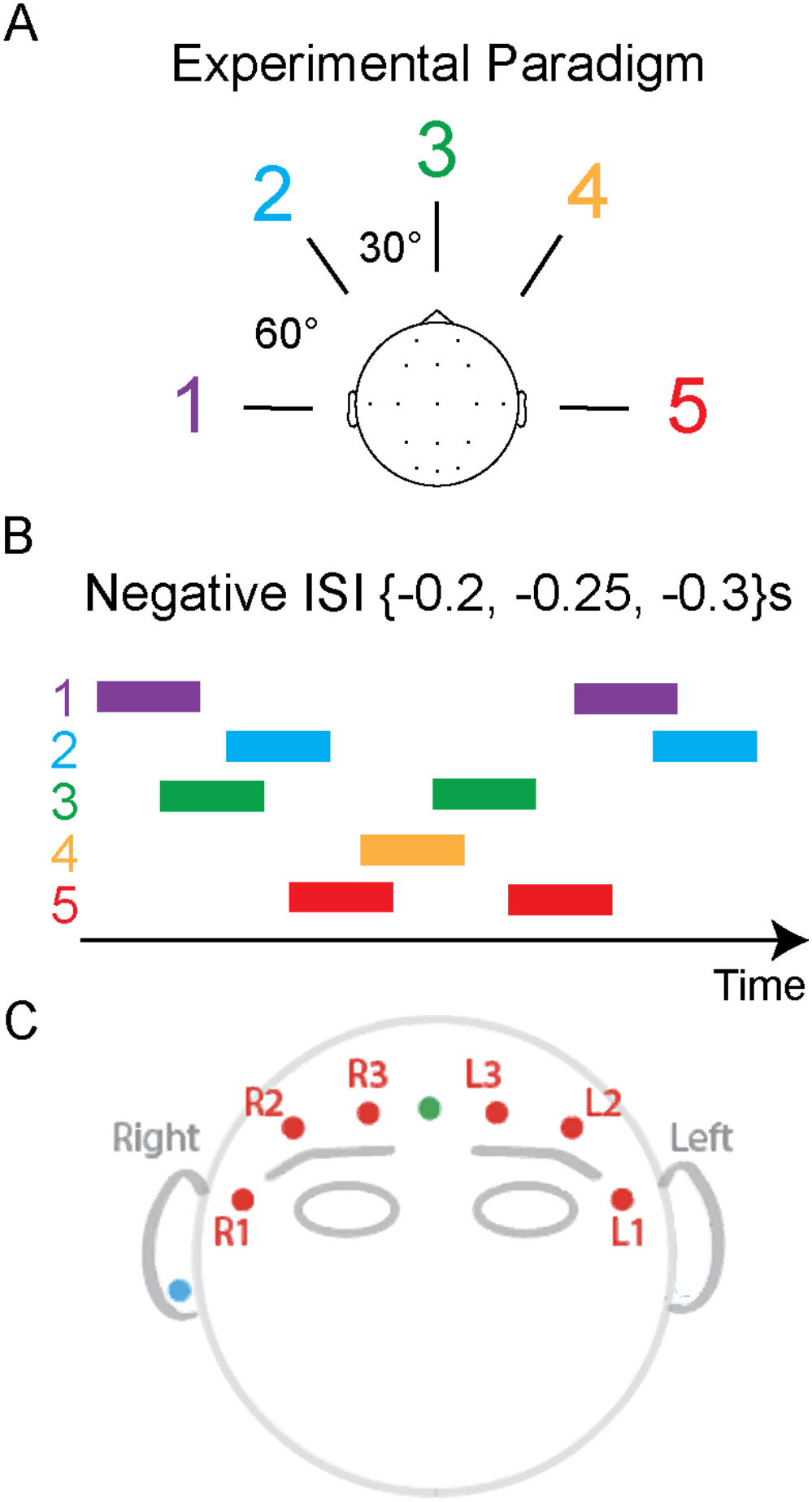
(A) Experimental design of AMUSE paradigm, which involves the presentation of five menu options presented from five spatial locations. (B) The beginning ∼3s of an example trial showing each sound, represented by colored rectangles, overlapped in time. (C) The locations of EOG electrodes.

Each trial began with a cue period during which a target cue word was displayed on a computer screen to indicate which of the five sounds the participant should pay attention to. This was followed by the presentation of the sound sequence. Each trial contained the presentation of five sounds (one target and four non-targets) four times, for a total of twenty sounds played per trial. Throughout each trial, consecutive sounds were overlapped with a random negative inter-stimulus interval (ISI) chosen from [-0.2, -0.25, -0.3] seconds (Fig. 1B). One block consisted of five trials (one for each target). The study included a total of five blocks.

### C. EOG data collection and preprocessing

The EOG data were collected using dry sticky electrodes recorded using a g.HIamp amplifier from g.tec (www.gtec.at). The data were recorded at a sampling frequency of 600 Hz, and a 58-62 Hz notch filter was applied to eliminate power line noise. Six EOG electrodes were placed around the participant’s eyes (Fig. 1C). The location of the reference electrode was on the left ear. Bipolar derivation was used to reference all the electrodes. Each electrode was then subtracted from each of the other electrodes, resulting in fifteen additional channels of bipolar derivations. Finally, the average value of the 0.5s pre-sequence period was subtracted from each channel for each sequence. Fig. 2A shows the data from one such bipolar derivation from an example subject over the course of one block. The y-axis shows the amplitude of the EOG when the eyes move to each of the five directions (5 colors) during 5 trials (in time). The amplitude of the EOG signal contains information about the direction of the target.

**Fig 2.**
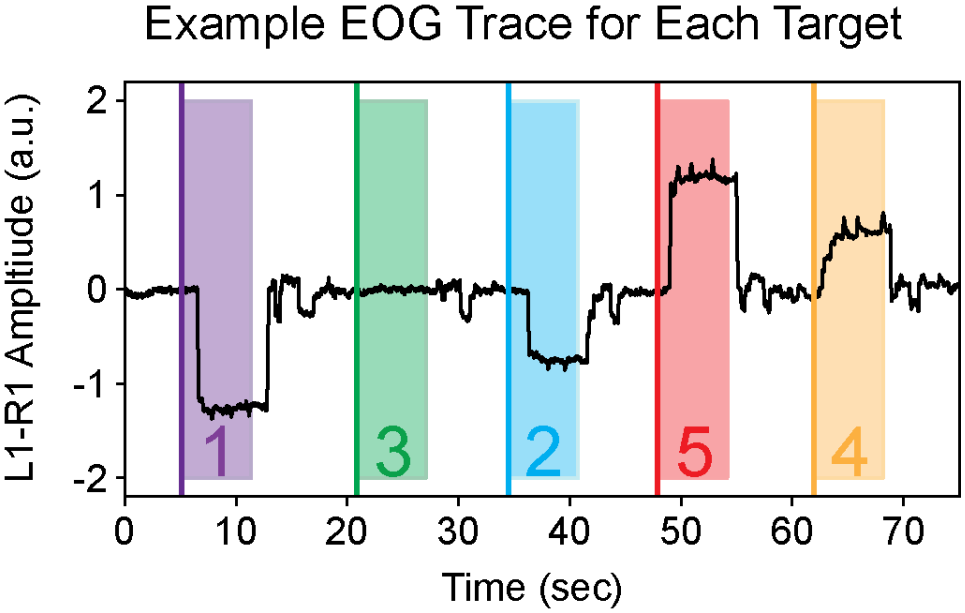
Raw bipolar L1-R1 EOG data from five trials (one block) of an example participant. Each shaded section and number corresponds to the time span and target word for each trial.

### D. Decoding Analysis

We trained linear discriminant analysis (LDA) classifiers using the EOG data from each participant in a leave-one-trial-out fashion. First, we classified each full sequence. Since an average of 1.75s is needed to present one full trial of all five options, the participant had to wait at most 1.75s to locate the target sound direction, after which the participant could turn their gaze. Therefore, we trained and tested the full-sequence EOG classifier on the data from all channels and time points starting from 1.75s until the end of the 6.25s sequence (4.5s). Next, we classified each moment in time. To determine the classification accuracy as a function of time over the course of a sequence, we trained and tested time-specific LDA classifiers on data from the previous 1.0s. This allowed us to determine how quickly we can achieve accurate decoding and what ITRs are possible when shortening the sequence duration. All modeling was done with scikit-learn [20].

## III. Results

### A. Full sequence EOG decoding

Fig. 3A shows the box plot of the average full-sequence decoding accuracy of the five menu options using LDA classification. The median accuracy is 80% across participants. We also show the average confusion matrix from all ten participants (Fig 3B) and the best-performing participant (Fig 3C). These results indicate that the proposed method can decode the attended target in the negative ISI AMUSE paradigm with an average accuracy of 76.5% across the ten participants and 100% for the best participant.

**Fig 3.**
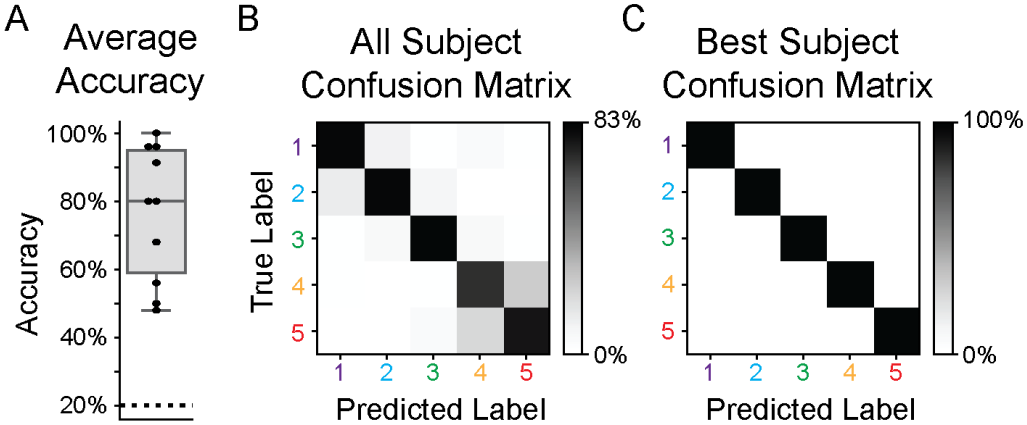
(A) Average decoding accuracy for the decoding of five menu options over 8 participants (B) Average confusion matrix over participants. (C) Confusion matrix for best participant.

### B. EOG decoding over time

Using time-specific LDA classifiers, we decoded the selected menu option over the course of each test sequence using EOG signals. Fig. 4A shows decoding accuracies for decoding 5 menu options over time. The results show that the accuracies peak at time t=1.95s and drop slightly as the sequence progresses. Fig 4B shows the average confusion matrix over subjects at t=1.95s when the average accuracy reaches its maximum of 82.7%.

**Fig 4.**
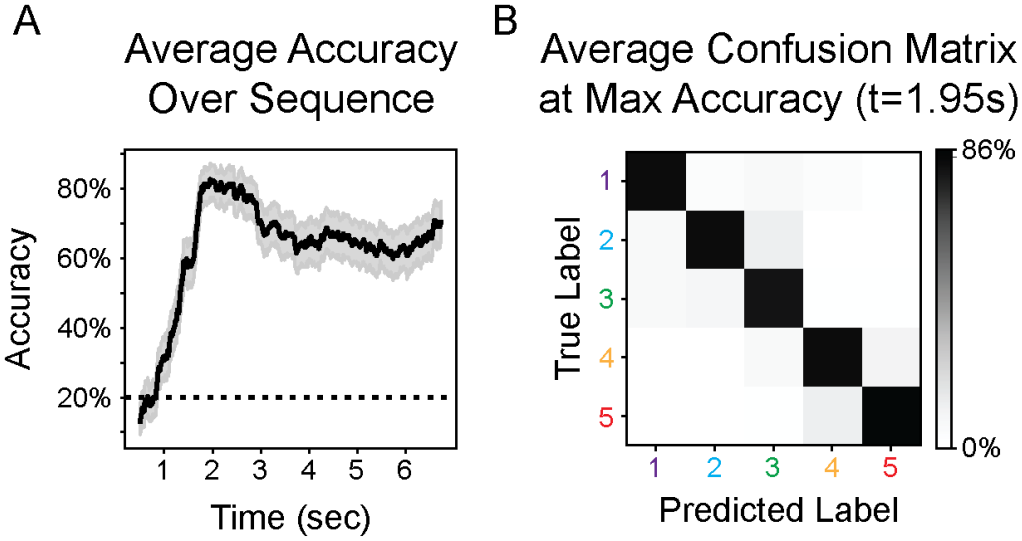
(A) Decoding 5 menu options over time, mean and standard error shown over 10 participants. (B) Average confusion matrix over participants at the time of maximum average accuracy of 82.7% (t=1.95s).

To further analyze these results, we calculated the ITRs based on the full sequence decoding (Fig 3A) and the accuracies obtained from decoding over time (Fig. 4A). At t=1.83s the average accuracy is 81.4% and results in a maximum average ITR of 43.6 bits/min. Table 1 shows the average and best participant ITRs for full sequence decoding and the maximum ITR when decoding over time.

**TABLE 1.** Information transfer rate (ITR)

| ***Decoding Time*** | ***Average ITR (bits/min)*** | ***Base Participant ITR (bits/min)*** |
|---|---|---|
| Full sequence | 13.4 | 22.3 |
| Peak ITR | 43.6 | 81.5 |

## IV. Discussion

In this investigation, we integrated a rapid AMUSE auditory HCI paradigm with EOG signals. This hybrid approach presents two main advantages. One, it eliminates the need for a display screen in HCI communication. Two, the integration of this paradigm with EOG signals demonstrates high performance (81.4%) in a brief decoding time of only 1.83 s, resulting in an average ITR of 43.6 bits/min. Notably, these performance metrics surpass the benchmarks set by the state-of-the-art EEG methods, whether using display screens (∼20 bits/min [15]) or engaging in non-display auditory HCI (∼10 bits/min [14]).

The methodologies outlined in this paper exhibit promising real-life applications for EOG-based HCI communication. These applications extend to scenarios where display-based interaction is inconvenient, such as driving, running, attending meetings, or using a device in public. In these contexts, the introduced methods prove effective for option selection, enabling control over a music player or the activation and deactivation of electronic devices. Moreover, their versatility is showcased when integrated with other inputs, such as managing incoming calls, offering the user choices like answering, rejecting, or selecting predefined settings. The study’s outcomes underscore the potential of achieving high performance (>80% for five classes) within a brief duration (1.83 seconds) through EOG-based gaze detection.

The accuracy of EOG is influenced by the sensor location, with different placements contributing differently to the decoding results. Our observations highlight that the bipolar signal from the L1 and R1 sensors (Fig. 2) contributed most to decoding for many participants; however, the contributions of EOG sensors varied across participants. Future investigations could explore the optimal sensor locations for EOG in different applications and form factors to enhance overall performance. We also note reduced accuracy for extremely peripheral targets, as it is difficult to orient gaze to such a large azimuth. Finally, incorporating feedback-based training and examining its impact on performance could offer valuable insights for further research.

## V. Conclusion

We employed a customized auditory AMUSE HCI paradigm for screen-free decoding of a selected option utilizing EOG-based eye-gaze classification. Through this paradigm, we attained an impressive ITR of 43.6 bits/min. This achievement significantly surpasses the current auditory and visual BCI paradigms which typically yield around ∼10 bits/min and ∼20 bits/min respectively.

